# DYT1 dystonia patient-derived fibroblasts have increased deformability and susceptibility to damage by mechanical forces

**DOI:** 10.1101/480186

**Authors:** Navjot Kaur Gill, Chau Ly, Paul H. Kim, Cosmo A. Saunders, Loren G. Fong, Stephen G. Young, G.W. Gant Luxton, Amy C. Rowat

## Abstract

DYT1 dystonia is a neurological movement disorder that is caused by a loss-of-function mutation in the *DYT1*/*TOR1A* gene, which encodes torsinA, the luminal ATPase-associated (AAA+) protein. TorsinA is required for the assembly of functional linker of nucleoskeleton and cytoskeleton (LINC) complexes, and consequently the mechanical integration of the nucleus and the cytoskeleton. Despite the potential implications of altered mechanobiology in dystonia pathogenesis, the role of torsinA in regulating cellular mechanical phenotype, or mechanotype, in DYT1 dystonia remains unknown. Here, we define the mechanotype of mouse fibroblasts lacking functional torsinA as well as human fibroblasts isolated from DYT1 dystonia patients. We find that the deletion of torsinA or the expression of torsinA containing the DYT1 dystonia-causing ΔE302/303 (ΔE) mutation results in a more deformable cellular mechanotype. We observe a similar increased deformability of mouse fibroblasts that lack lamina-associated polypeptide 1 (LAP1), which interacts with and stimulates the ATPase activity of torsinA *in vitro*; as well as with depletion of the LINC complex proteins, Sad1/UNC-84 (SUN)1 and SUN2, lamin A/C, or lamin B1. Moreover, we report that DYT1 dystonia patient-derived fibroblasts are more compliant than fibroblasts isolated from unafflicted individuals. DYT1 fibroblasts also exhibit increased nuclear strain and decreased viability following mechanical stretch. Taken together, our results support a model where the physical connectivity between the cytoskeleton and nucleus contributes to cellular mechanotype. These findings establish the foundation for future mechanistic studies to understand how altered cellular mechanotype may contribute to DYT1 dystonia pathogenesis; this may be particularly relevant in the context of how neurons sense and respond to mechanical forces during traumatic brain injury, which is known to be a major cause of acquired dystonia.

## 1 Introduction

Dystonia is a debilitating ‘hyperkinetic’ neurological movement disorder, which is the third most common movement disorder worldwide behind essential tremor and Parkinson’s disease (Fahn, 1988; Fahn et al., 1988; Geyer and Bressman, 2006; Defazio et al., 2007). Dystonia is characterized by involuntary sustained or intermittent muscle contractions resulting in abnormal repetitive movements and/or postures (Fahn, 1988; Albanese et al., 2013). While there are multiple treatment options to manage dystonia—such as botulinum toxin injection, oral medications, and deep brain stimulation—no curative therapies are available (Albanese et al., 2013). If we could fully define the mechanisms of disease pathogenesis, this would enable the development of effective targeted treatment strategies for dystonia patients.

Dystonia can be acquired as a result of traumatic brain injury, central nervous system infection, or environmental toxins (Albanese et al., 2013; Albanese et al., 2018). This neurological disorder can also be inherited: the most prevalent and severe inherited dystonia (Weisheit et al., 2018), DYT1 dystonia, is caused by a loss-of-function mutation in the *DYT1*/*TOR1A* gene that deletes a single glutamic acid residue (ΔE302/303, or ΔE) from the encoded torsinA protein (Ozelius et al., 1997). TorsinA is a ATPase-associated with various cellular activities (AAA+) protein, which resides within the lumen of the endoplasmic reticulum lumen and the contiguous perinuclear space of the nuclear envelope (Goodchild and Dauer, 2004; Naismith et al., 2004). AAA^+^ proteins typically function as ATP-dependent molecular chaperones that structurally remodel their protein substrates (Hanson and Whiteheart, 2005). While the substrate(s) remodeled by torsinA are unknown, torsinA is thought to function within the nuclear envelope where its ATPase activity is stimulated by its membrane-spanning co-factors: lamina-associated polypeptide 1 (LAP1) and luminal domain-like LAP1 (LULL1) (Laudermilch and Schlieker, 2016). While the ΔE mutation impairs the ability of torsinA to interact with or be stimulated by either LAP1 or LULL1 (Naismith et al., 2009; Zhao et al., 2013), a mechanistic understanding of how the ΔE mutation drives DYT1 dystonia pathogenesis at the cellular level remains unclear.

We recently identified torsinA and LAP1 as mediators of the assembly of functional linker of nucleoskeleton and cytoskeleton (LINC complexes) (Saunders and Luxton, 2016; Saunders et al., 2017), which are evolutionarily conserved nuclear envelope-spanning molecular bridges that mechanically integrate the nucleus and the cytoskeleton (Kaminski et al., 2014; Chang et al., 2015b). LINC complexes are composed of the outer and inner nuclear membrane nesprin and SUN proteins: nesprins interact with the cytoskeleton in the cytoplasm and SUN proteins in the perinuclear space, whereas SUN proteins interact with A-type lamins and chromatin-binding proteins in the nucleoplasm (Crisp et al., 2006; Wilson and Berk, 2010; Chang et al., 2015a). Our previous work demonstrated that torsinA and LAP1 are required for the assembly of transmembrane actin-associated nuclear (TAN) lines (Saunders et al., 2017), which are linear arrays of LINC complexes containing the actin-binding nesprin-2Giant (nesprin-2G) and SUN2 that harness the forces generated by the retrograde flow of perinuclear actin cables to move the nucleus towards the rear of migrating fibroblasts and myoblasts (Luxton et al., 2010; Luxton et al., 2011; Chang et al., 2015a). Consistent with these findings, DYT1 dystonia patient-derived fibroblasts and fibroblasts isolated from mouse models of DYT1 dystonia exhibit reduced motility *in vitro* (Nery et al., 2008; Nery et al., 2014). Moreover, the migration of torsinA-null neurons in the dorsal forebrain of mouse embryos *in vivo* show impaired migration (McCarthy et al., 2012; Nery et al., 2014). Since intracellular force generation is critical for cell motility, and regulated by shared mediators of mechanotype, these results suggest that DYT1 dystonia may be characterized by defective mechanobiology.

Here we test the hypothesis that torsinA regulates cellular mechanical phenotype, or mechanotype, which describes how cells deform in response to mechanical stresses. Cellular mechanotype is critical for mechanotransduction, whereby cells translate mechanical stimuli from their environment into biochemical signals and altered gene expression (Tyler, 2012; Franze, 2013). The ability of cells to withstand physical forces is also critical for their survival (Hsieh and Nguyen, 2005). For example, the external stresses of traumatic brain injury result in cell death (Raghupathi, 2004; Stoica and Faden, 2010; Hiebert et al., 2015; Ganos et al., 2016). Damage to cells can also occur during migration through narrow constrictions, including nuclear rupture, DNA damage, and ultimately cell death (Harada et al., 2014; Denais et al., 2016; Raab et al., 2016; Irianto et al., 2017). The damaging effects of such large cellular deformations depend on levels of A-type nuclear lamins, which are critical regulators of nuclear and cellular mechanotype (Lammerding et al., 2004; Swift et al., 2013; Stephens et al., 2017). The depletion of other proteins that associate with nuclear lamins, such as the inner nuclear membrane protein emerin, also result in reduced mechanical stability of the nuclear envelope (Rowat et al., 2006; Reis-Sobreiro et al., 2018) and increased nuclear deformability in response to strain (Lammerding et al., 2005). The nuclear lamina interacts with chromatin, which can also contribute to nuclear mechanical properties (Pajerowski et al., 2007; Chalut et al., 2012; Schreiner et al., 2015; Stephens et al., 2017). In addition, nuclear lamins associate with the LINC complex, which is an important mediator of the transmission of physical forces generated from the cytoskeleton to the nucleus (Stewart-Hutchinson et al., 2008; Lombardi et al., 2011; Spagnol and Dahl, 2014). Given that torsinA is required for the assembly of functional LINC complexes (Saunders et al., 2017), we speculated that DYT1 dystonia cells may exhibit altered mechanotype, which could contribute to the pathogenesis of DYT1 dystonia.

Here we show that cellular mechanotype is altered due to the expression of torsinA containing the DYT1 dystonia-causing ΔE mutation, the deletion of torsinA, or the disruption of functional LINC complexes. We use fibroblasts as a model system, which have been used successfully to model human neurological disorders, including dystonia (Connolly, 1998; Hewett et al., 2006; Auburger et al., 2012; Burbulla and Kruger, 2012; Wray et al., 2012; Nery et al., 2014). We find that mouse embryonic fibroblasts (MEFs) derived from torsinA-or LAP1-knockout as well as ΔE-knock-in mice are more deformable than control fibroblasts. We observe a similar, more compliant mechanotype in MEFs lacking functional LINC complexes as well as A-or B-type lamins. Furthermore, we find that fibroblasts isolated from DYT1 dystonia patients are more deformable than normal fibroblasts. Interestingly we observe that DYT1 fibroblasts exhibit nuclei with greater strain and decreased cell viability following mechanical stretching. Collectively these findings establish altered cellular mechanotype as a potential biomarker of DYT1 dystonia, which will guide future studies designed to better understand dystonia pathogenesis and to identify novel therapeutic targets for treatment of this debilitating disease.

## 2 Materials and methods

### 2.1 Cells

Parental NIH3T3 fibroblasts were cultured in L-glutamine-, glucose-, and sodium pyruvate-containing Dulbecco’s modified Eagle’s media (DMEM) (ThermoFisher Scientific, Waltham, MA) supplemented with 10% bovine calf serum (BCS) (Gemini Bio-Products, West Sacramento, CA). NIH3T3 fibroblasts stably expressing wild type (WT), or mutant (E171Q or ΔE) versions of torsinA containing EGFP inserted after its signal sequence (SS) were created as follows: the open reading frame encoding SS-EGFP-torsinA was amplified by PCR from the previously described SS-EGFP-torsinA WT, SS-EGFP-torsinA E171Q and SS-EGFP-torsinA ΔE constructs (Saunders et al., 2017) using the primers SS-EGFP-F (5’-GGGCGCCTCGAGATGAAGCTGGGCCGGG-3’) and SS-EGFP-torsinA-R (5’-GCGCCCGAATTCTCAATCATCGTAGTAATAATCTAACTTGGTG-3’), which contain 5’ *Xho*I and *Eco*RI cut sites, respectively. The resulting PCR products were each purified using the Wizard SV Gel and PCR Clean-Up System (Promega, Madison, WI) and then subcloned into the cytomegalovirus immediate-early expression cassette of the pLPCX retroviral vector (Takara Bio USA, Inc., Mountain View, CA). Phusion DNA polymerase and T4 DNA ligase were purchased from New England Biolabs (Ipswich, MA). Restriction enzymes were either purchased from New England Biolabs or Promega. The resultant pLPCX cDNA constructs and pVSV-G (Takara Bio USA, Inc.) were purified using the GeneJet Plasmid Midiprep Kit (ThermoFisher Scientific) and cotransfected into the gp293 retroviral packaging cell line (Takara Bio USA, Inc.); the subsequent isolation of retroviral particles was performed as recommended by the manufacturer. NIH3T3 fibroblasts were transduced with the resultant retrovirus and selected with 2 µg/mL puromycin (Thermo Fisher Scientific). Individual clones of the resultant cell lines were isolated using limiting dilution and maintained with 2 µg/mL of puromycin. Mouse embryonic fibroblasts (MEF) lacking Tor1a, Tor1aip1, and SUN1/2 as well as lamin A and B1 were derived from mice as previously described (Kim et al., 2010; Jung et al., 2014). All mouse MEFs used in this study were grown in DMEM with 15% BCS. Human fibroblasts (GM00023, GM00024, GM02912, GM02551, GM03208, GM03211, GM03221, and GM02304) were purchased from the Coriell Institute and cultured following vendor’s instructions (Camden, NJ): GM00023, GM03208, GM03211, GM03221, GM02551 and GM02304 were grown in DMEM containing 15% fetal bovine serum (FBS) (Gemini Bio-Products, West Sacramento, CA). GM00024 were grown in DMEM supplemented with 10% FBS. GM02912 were grown in 20% FBS-containing Ham’s F12 media supplemented with 2 mM L-glutamine (Sigma-Aldrich, St. Louis, MO).

### 2.2 Parallel microfiltration

Prior to filtration measurements, cells were washed with 1x DNase-, RNase-& Protease-free phosphate-buffered saline purchased from Mediatech (Manassas, VA), treated with trypsin (VWR, Visalia, CA), and resuspended in fresh medium to a density of 0.5 × 10^6^ cells/mL. Cell suspensions were then passed through a 35 µm cell strainer (BD Falcon, San Jose, CA) prior to each filtration measurement. Next, 350 µL of each cell suspension was loaded into each well of a 96-well loading plate (Greiner BioOne, Monroe, NC). The number and size distribution of cells in each well was quantified using an automated cell counter (TC20, BioRad Laboratories, Hercules, CA). Finally, a defined amount of air pressure, which was monitored using a 0 −100 kPa pressure gauge (Noshok Inc., Berea, OH), was applied to the 96-well plate outfitted with a custom pressure chamber (Qi et al., 2015; Gill et al., 2017). To quantify retention volumes following filtration, we measured the absorbance at 560 nm of the phenol red-containing cell medium using a plate reader (Infinite M1000, Tecan Group Ltd., Männedorf, Switzerland).

### 2.3 q-DC

Standard soft lithography methods were used to fabricate the microfluidic devices for q-DC experiments. Briefly, a 10:1 w/w base to crosslinker ratio of polydimethylsiloxane (PDMS) was poured onto a previously described master wafer (Saunders et al., 2017). The device was subsequently bonded to a No. 1.5 glass coverslip (Thermo Fisher) using plasma treatment (Plasma Etch, Carson City, NV). Within 24 h of device fabrication, suspensions of 2 × 10^6^ cells/mL were driven through constrictions of 9 μm (width) x 10 μm (height) by applying 55 kPa of air pressure across the device. We captured images of cell shape during transit through the 9 µm gaps on the millisecond timescale using a CMOS camera with a capture rate of 1600 frames/s (Vision Research, Wayne, NJ) mounted on an inverted Axiovert microscope (Zeiss, Oberkochen, Germany) equipped with a 20x/0.4NA objective (LD Achroplan 20x/0.4NA objective Korr Ph2, Zeiss) and light source (Osram Halogen Optic Lamp 100 W, 12 V). We used custom MATLAB (MathWorks, Natick, MA) code (https://github.com/knybe/RowatLab-DC-Analysis) to analyse the time-dependent shape and position changes of individual cells (Nyberg et al., 2016). To determine the mechanical stresses applied to individual cells, we used agarose calibration particles that were fabricated using oil-in-water emulsions as previously described (Nyberg et al., 2017). Stress-strain curves were obtained for single cells and a power-law rheology model was subsequently fitted to the data to compute the elastic modulus and fluidity of the cells.

### 2.4 Epifluorescence microscopy

To image cell and nuclear morphology, cells grown on No. 1.5 coverslips were labeled with 5 µM Calcein-AM and 0.2 µg/mL Hoechst 33342 (ThermoFischer Scientific). For cell viability measurements, cells were stained with 50 µg/mL propidium iodide (Thermo Fisher Scientific). Images of fluorescently-labeled cells were acquired using a Zeiss Axio Observer A.1 microscope equipped with a 10x/0.3 NA EC Plan-Neofluar Ph1 M27 objective, 20x/0.8 NA Plan-Apochromat M27 objective, HBO 103W/2 mercury vapor short-arc lamp light source, BP 470/20 excitation filter, BP 505-530 emission filter, and FT 495 beam splitter. ImageJ (Bethesda, MA) was used to quantify cell and nuclear size and shape parameters from the acquired images.

### 2.5 Cell stretching

To subject cells to external mechanical stresses, we used a custom-built cell stretching apparatus (Kim et al., 2018). We prepared elastic PDMS membranes as previously described (Kim et al., 2018). Cells were resuspended in tissue culture media at a concentration of 5 x 10^5^ cells/mL and then added to individual PDMS strips and incubated for 24 h at 37 °C in cell culture incubator. To quantify nuclear strain, the membranes with cells adhered were stretched by 2 mm (5% of the total length of the membrane) while submerged in cell culture media for 5 min prior to imaging. To determine effects of repetitive stretch on cell viability and adhesion, membranes were stretched by 2 mm at 0.5 Hz for 24 h at 37 °C. After 24 h, membrane-adhered cells were stained with fluorescent dyes and imaged as described above. To quantify the number of cells attached to the membranes after stretching, we prepared lysates of the adherent cells using a solution of 0.1 N NaOH (Sigma Aldrich) and measured the total protein content of the lysates using the BioRad Laboratories D-C protein assay kit.

### 2.6 Statistical methods

To determine statistical significance of data that exhibited non-parametric distributions including transit time, apparent cell elastic modulus, cell size, cell and nuclear shape, and nuclear strain, we used the Mann-Whitney U test. All other results are expressed as mean ± standard deviation and Student’s t-test method was used to analyze significance and to obtain p-values.

## 3 Results

### 3.1 TorsinA and LAP1 contribute to fibroblast mechanotype

To begin to determine if DYT1 dystonia is associated with altered cellular mechanotype, we performed parallel microfiltration (PMF) (Qi et al., 2015; Gill et al., 2017) of NIH3T3 fibroblasts that have inhibited function of torsinA. In PMF, cell suspensions are filtered through porous membranes by applying air pressure. Cells that occlude the micron-scale pores due to their stiffness and/or size block the fluid flow, reducing filtrate volume, and increasing the volume of fluid that is retained in the top well, which we report as % retention (**Figure 1A**). To manipulate torsinA function, we generated lentivirus-transduced NIH3T3 fibroblasts that stably express previously described cDNA constructs encoding wild type (WT) or mutant (E171Q or ΔE) torsinA (Saunders et al., 2017). Since neither SS-EGFP-torsinA^E171Q^ nor SS-EGFP-torsinA^ΔE^ were able to rescue the rearward nuclear positioning defect in mouse embryonic fibroblasts (MEFs) isolated from torsinA-knockout (*Tor1a*^-/-^) mice (Goodchild et al., 2005; Saunders et al., 2017), we rationalized that these constructs would act as dominant negative inhibitors of torsinA function in NIH3T3 fibroblasts.

**Figure 1.**
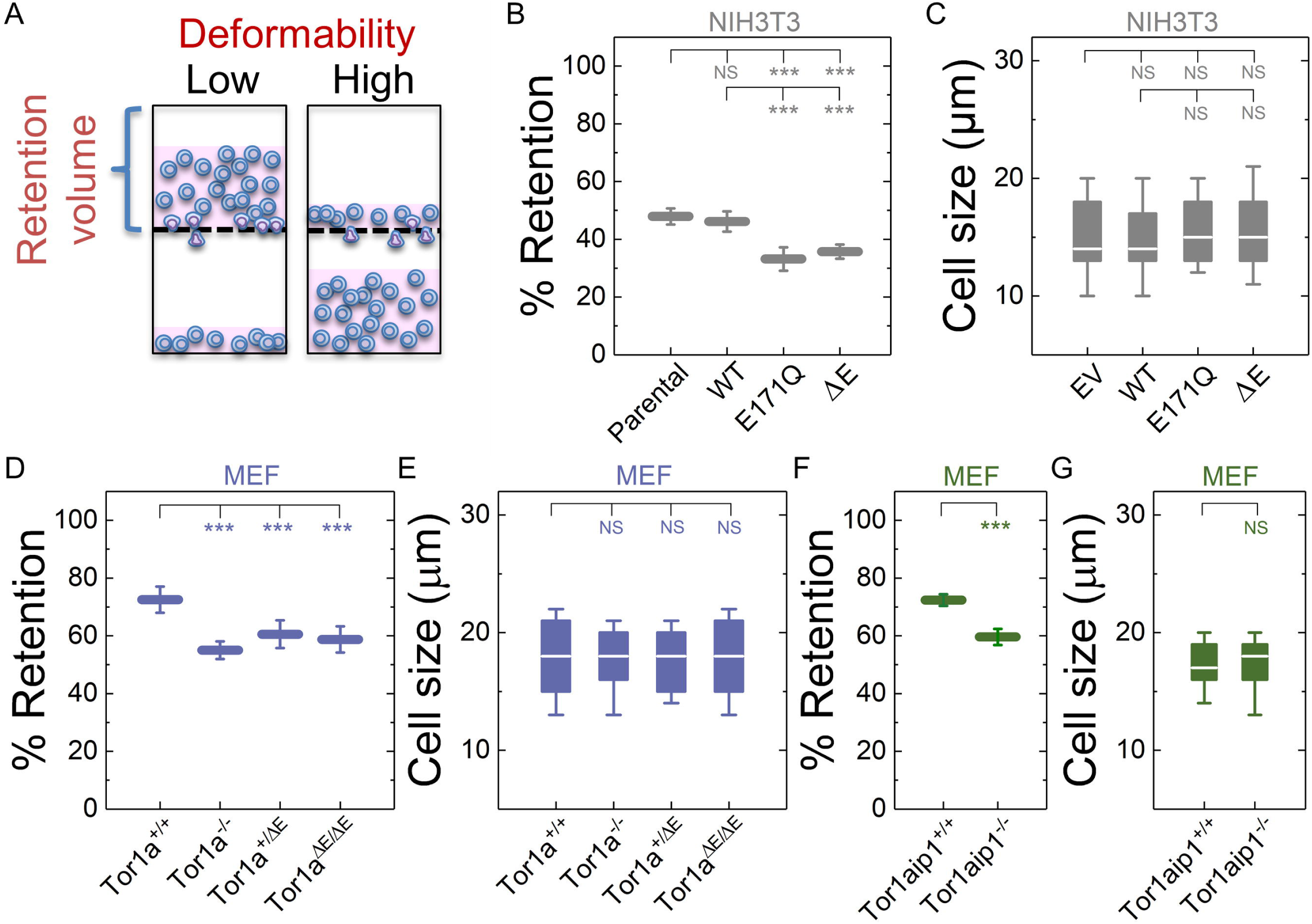
Fibroblasts lacking functional torsinA or LAP1 have increased deformability. (**A**) Schematic illustration of PMF. Less deformable cells tend to occlude pores, which blocks fluid flow and results in an increased volume of fluid that is retained above the membrane (% retention). (**B**) PMF measurements of NIH3T3 fibroblasts expressing the indicated SS-EGFP-tagged torsinA constructs. (**D, F**) PMF measurements of the indicated MEF lines. PMF conditions: 10 µm pore size, 2.1 kPa for 50 s. Each data point represents mean ± standard deviation (SD). Statistical significance was determined using the Student’s t-test. (**C, E, G**) Cell size measurements. Boxplots show 25^th^ and 75^th^ percentiles; line shows median; and whiskers denote 10^th^ and 90^th^ percentiles. All data were obtained from three independent experiments. Statistical significance was determined using the Mann-Whitney U test. *** p < 0.001; ** p < 0.01; not significant (NS) p > 0.05.

Consistent with this prediction, NIH3T3 fibroblasts expressing either SS-EGFP-torsinA^E171Q^ or SS-EGFP-torsinA^ΔE^ exhibited significantly lower % retention than parental non-transduced or SS-EGFP-torsinA^WT^ transduced NIH3T3 fibroblasts (**Figure 1B**). While cell size can impact filtration, we observed no significant differences in size distributions among these cell lines (**Figure 1C**), suggesting that their altered filtration is due to differences in their cellular deformability. These data suggest that torsinA regulates the mechanotype of NIH3T3 fibroblasts. To further investigate the relationship between torsinA and cellular mechanotype, we next performed PMF experiments on previously characterized *Tor1a*^+/+^ and *Tor1a*^-/-^ MEFs (Saunders et al., 2017). We found that % retention of the *Tor1a*^+/+^ MEFs was significantly larger than the % retention of the *Tor1a*^-/-^ MEFs (**Figure 1D**). Consistent with these findings, we observed that MEFs isolated from heterozygous (*Tor1a*^+/ΔE^) or homozygous (*Tor1a*^ΔE/ΔE^) ΔE-knock-in mice (Goodchild et al., 2005) had a significantly lower % retention than *Tor1a*^+/+^ MEFs (**Figure 1D**). Moreover, the % retention measured for MEFs derived from LAP1-knockout (*Tor1aip1*^-/-^) mice was also significantly lower than control *Tor1aip1*^+/+^ MEFs (**Figure 1F**). We confirmed that these observed changes in % retention were not due to significant differences in cell size distributions (**Figures 1E, G**). Because the interaction between torsinA and the luminal domain of LAP1 stimulates its ATPase activity *in vitro* (Zhao et al., 2013) and the ΔE mutation impairs the ability of torsinA to interact with LAP1 (Naismith et al., 2009), these results suggest that the interaction between torsinA and LAP1 may contribute to fibroblast mechanotype. In addition, LAP1 is critical for nuclear envelope structure (Santos et al., 2015) and interacts with nuclear lamins (Foisner and Gerace, 1993; Serrano et al., 2016), which are major determinants of cellular mechanotype (Houben et al., 2007). Thus, the loss of interaction between torsinA and LAP1 caused by the ΔE mutation may also contribute to the more compliant mechanotype that we observe in LAP1-knockout fibroblasts.

### 3.2 Components of the LINC complex mediate cellular mechanotype

We previously showed that torsinA and LAP1 are both required for nuclear-cytoskeletal coupling through SUN2-containing LINC complexes (Saunders et al., 2017). Thus, we next asked whether or not MEFs isolated from SUN2-knockout (*SUN2*^-/-^) mice (Lei et al., 2009) exhibit a similar decreased filtration as the *Tor1a*^-/-^ and *Tor1aip1*^-/-^ MEFs. We found that *SUN2*^-/-^ MEFs had a reduced % retention relative to control (*SUN1/2*^+/+^) MEFs (**Figure 2A**). Since torsinA has also been proposed to interact with and regulate SUN1-containing LINC complexes (Jungwirth et al., 2011; Pappas et al., 2018), we also performed PMF on MEFs derived from SUN1/2-double knockout (*SUN1/2*^-/-^) mice (Lei et al., 2009) and found that *SUN1/2*^-/-^ MEFs had reduced retention compared to both *SUN2*^-/-^ and *SUN1/2*^+/+^ MEFs (**Figure 2A**). We confirmed that these changes in % retention were not due to significant differences in cell size distributions (**Figure 2B**).

**Figure 2.**
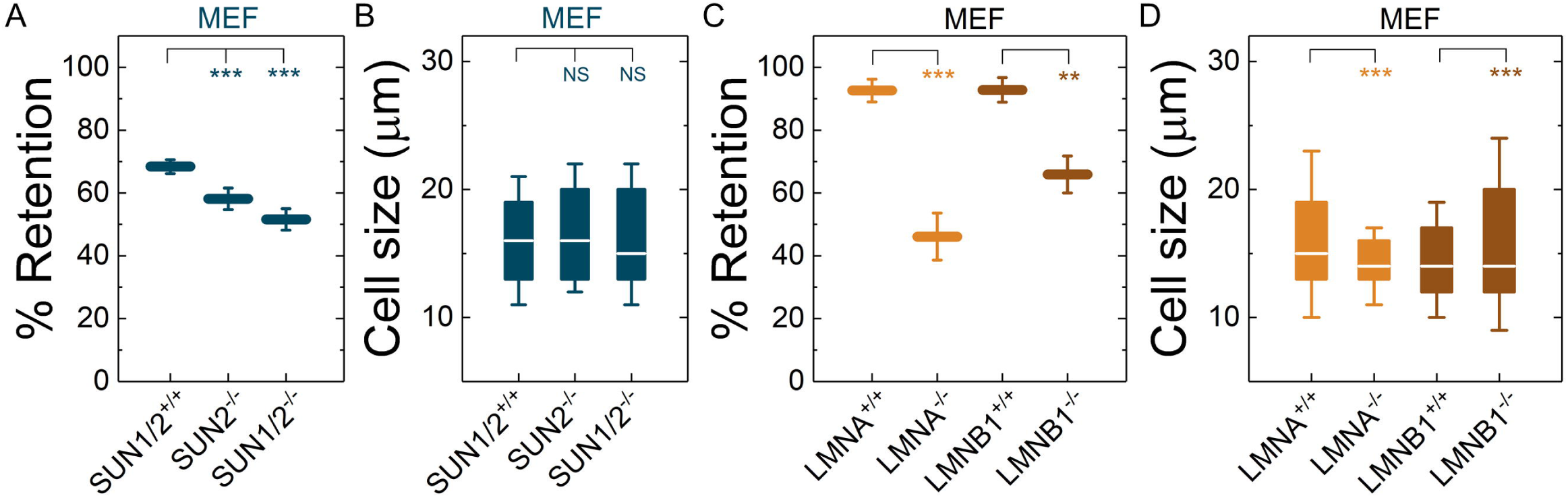
Fibroblasts lacking LINC complexes, A-type lamins, or lamin B1 have increased deformability. PMF measurements of the indicated MEF lines. (**A**) PMF conditions: 10 µm pore membrane, 2.1 kPa for 50 s. (**C**) PMF conditions: 10 µm pore membrane, 2.1 kPa for 40 s. Each data point represents mean ± SD. Statistical significance was determined using the Student’s t-test. (**B, D**) Cell size data. Boxplots show 25^th^ and 75^th^ percentiles; line shows median; whiskers denote 10^th^ and 90^th^ percentiles. Statistical significance was determined using the Mann-Whitney U test. All data were obtained from three independent experiments. *** p < 0.001; ** p < 0.01; not significant (NS) p > 0.05.

We next investigated the effects of A-type lamins on cell filtration, as they are established regulators of cellular mechanotype (Lammerding et al., 2004; Swift et al., 2013) and directly interact with LAP1 (Foisner and Gerace, 1993; Serrano et al., 2016) and SUN1/2 (Chang et al., 2015b). We found that MEFs isolated from lamin A/C-knock-out (*LMNA*^-/-^) mice exhibited a reduced % retention relative to MEFs isolated from control mice (*LMNA*^+/+^) (**Figure 2C**). The increased deformability of the *LMNA*^-/-^ MEFs that we observed is consistent with previous studies from our laboratory and others, which show that A-type lamins determine the ability of cells to deform through micron-scale pores, both during passive deformation driven by applied pressure (timescale ∼ seconds) and active migration (timescale ∼ hours) (Rowat et al., 2013; Denais et al., 2016; Irianto et al., 2017). In addition to A-type nuclear lamins, cells express the B-type nuclear lamins, lamin B1 and lamin B2 (Dittmer and Misteli, 2011; Reddy and Comai, 2016). Lamin B1 interacts with SUN1 (Nishioka et al., 2016) and LAP1 (Maison et al., 1997) and is required for proper nuclear-cytoskeletal coupling (Ji et al., 2007). We found that MEFs isolated from lamin-B1-knockout mice (*LMNB1*^-/-^) (Vergnes et al., 2004) had reduced % retention compared to MEFs isolated from control mice (*LMNB1*^+/+^) (**Figure 2C**), suggesting that lamin B1 also contributes to cellular mechanotype. These findings are in agreement with previous findings that lamin B1 is a determinant of nuclear shape and stiffness (Coffinier et al., 2011; Ferrera et al., 2014). While we observed differences in cell size distributions between *LMNA*^-/-^ and *LMNA*^+/+^ MEFs as well as between *LMNB1*^-/-^ and *LMNB1*^+/+^ MEFs, we did not observe that cells with larger median cell size had increased % retention (**Figures 2C, D**). Taken together, these results suggest that nuclear lamins, torsinA, LAP1, and LINC complexes are important mediators of cellular mechanotype. These results are consistent with a model where cells are more deformable when the mechanical integration of the nucleus and the cytoskeleton is perturbed.

### 3.3 Dystonia patient-derived fibroblasts are more deformable than controls

Having established that the DYT1 dystonia-causing ΔE mutation in torsinA makes cells more deformable and that torsinA and LINC complex-associated proteins are important determinants of cellular mechanotype, we next tested if fibroblasts isolated from DYT1 dystonia patients display defects in cellular mechanotype. We performed PMF on a panel of age-matched human fibroblasts isolated from normal control (GM00023, GM00024, GM02912) and DYT1 patients (GM03211, GM03221, GM02304). We found that DYT1 patient-derived fibroblasts had consistently lower % retention compared to fibroblasts isolated from unafflicted controls (**Figure 3A**). We did not observe any significant effects of cell size on retention measurements of these human fibroblasts. While there were some differences in cell size distributions across DYT1 fibroblast lines (**Figure 3B**), we consistently observed decreased % retention of the DYT1 dystonia patient-derived fibroblasts, even for the GM02304 line that has a larger median cell size (**Figure 3A**). These findings suggest that fibroblasts isolated from DYT1 dystonia patients more readily deform through micron-scale pores.

**Figure 3.**
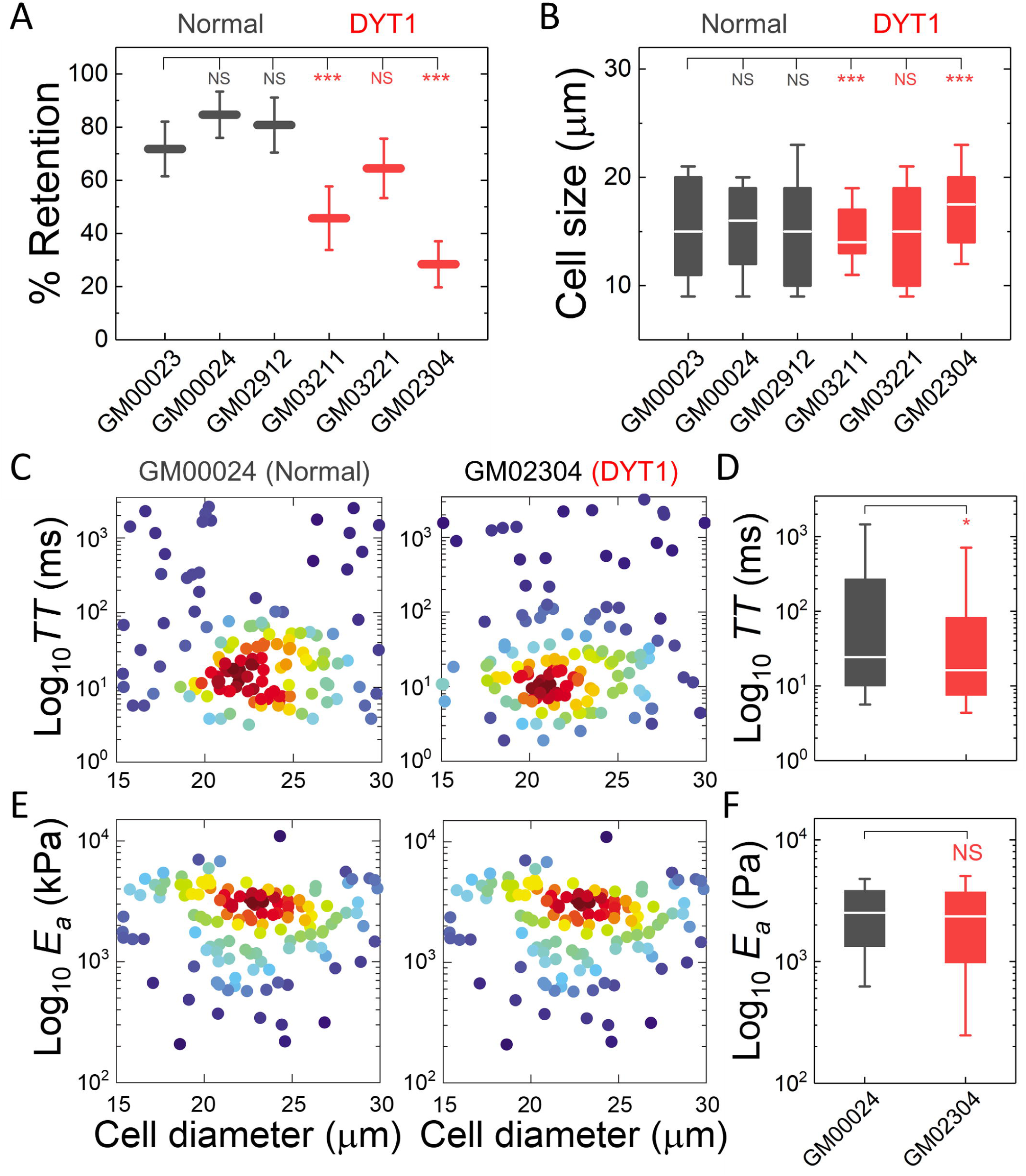
Dystonia patient-derived fibroblasts have increased deformability. (**A**) PMF measurements of normal and DYT1 patient-derived fibroblast lines. PMF conditions: 10 µm pore membrane, 1.4 kPa for 50 s. Each data point represents mean ± SD. Statistical significance was determined using the Student’s t-test. (**B**) Cell size distributions. Boxplots show 25^th^ and 75^th^ percentiles; line shows median; whiskers denote 10^th^ and 90^th^ percentiles. (**C, E**) Density scatter plots for *TT* and *E*_*a*_ measurements determined by q-DC. Each dot represents a single cell. N > 190 per sample. (**D, F**) *TT* and *E*_*a*_ measurement boxplots show 25^th^ and 75^th^ percentiles; line shows median; whiskers denote 10^th^ and 90^th^ percentiles. Statistical significance was determined using Mann-Whitney U test. All data were obtained over three independent experiments. *** p < 0.001; ** p < 0.01; not significant (NS) p > 0.05.

To validate the altered mechanotype of the DYT1 patient-derived fibroblasts (GM02304) versus control cells (GM00024), we used quantitative deformability cytometry (q-DC) (Nyberg et al., 2017). q-DC is a microfluidic method that enables single-cell measurements of transit time (*TT*), which is the time that it takes a cell to transit into the micron-scale constriction of a microfluidic device in response to applied pressure, and apparent elastic modulus (*E*_*a*_). We found that DYT1 fibroblasts had reduced median *TT* relative to normal fibroblasts (median *TT*_GM02304_ = 16.2 ms versus *TT*_GM00024_ = 24.4 ms, p = 1.5 x 10^-2^). Since *TT* tends to be shorter for more compliant cells with reduced elastic modulus (Nyberg et al., 2017), these findings corroborate the increased deformability of DYT1 fibroblasts (**Figures 3C, D**). q-DC measurements can also be impacted by cell size, but we found no significant correlations of q-DC measurements with cell diameter (*d*) by linear regression analysis (Pearson’s r_GM02304_*TT* vs *d*_ = 0.0, Pearson’s r_GM00024_*TT* vs *d*_ = −0.1), suggesting that these observations of the altered DYT1 dystonia-derived fibroblast mechanotype does not depend on cell size. Using power law rheology to extract measurements of *E*_*a*_ (Nyberg et al., 2017), we found that DYT1 patient-derived fibroblasts have reduced median *E*_*a*_ compared to controls, although the reduction was not statistically significant (**Figures 3E, F**). Collectively, our PMF and q-DC measurements indicate that DYT1 dystonia patient-derived fibroblasts are more deformable than control fibroblasts.

### 3.4 DYT1 dystonia patient-derived fibroblasts display altered nuclear morphology

Cellular and nuclear shape reflect a balance between cell-matrix adhesion, cellular force generation, mechanical stability of the cellular cortex and nuclear envelope, as well as nuclear-cytoskeletal connectivity (Dahl et al., 2004; Dahl et al., 2008; Rowat et al., 2008; Kim et al., 2015; Murrell et al., 2015). Since the ΔE mutation in torsin A alters the mechanical integration of the nucleus and the cytoskeleton via the LINC complex (Nery et al., 2008; Jungwirth et al., 2011; Saunders et al., 2017), and we observed differences in cellular deformability between the DYT1 dystonia patient-derived fibroblasts and control fibroblasts, we hypothesized that fibroblasts isolated from DYT1 dystonia patients may exhibit altered cellular size as well as nuclear size and shape. To characterize these features, we performed quantitative image analysis of cells with labeled cytoplasm (Calcein AM) and nuclei (Hoechst) using epifluorescence microscopy. We first measured the area of adhered cells, which indicates the degree of cell spreading. Based on our observations of the more compliant mechanotype of DYT1 dystonia patient-derived fibroblasts and a previous report of the altered adhesion of these cells (Hewett et al., 2006), we hypothesized that DYT1 fibroblasts may have reduced spread area. However, we found no observable differences in cellular area between DYT1 and normal fibroblasts (**Supplementary Figure 1A**). Since intracellular forces pulling on the nucleus can result in an increased nuclear area (Iyer et al., 2012), we next measured the projected area of nuclei in these cells, but found no significant differences between fibroblasts isolated from DYT1 dystonia patients or controls (**Supplementary Figure 1B**). Cell-to-nuclear size ratio was also similar across cell types, indicating that nuclear and cellular size scale similarly in DYT1 dystonia patient-derived fibroblasts and control cells (**Supplementary Figure 1C**).

We next investigated nuclear shape, which is impacted by cytoskeletal-generated forces as well as the inherent mechanical stability of the nuclear envelope (Rowat et al., 2008; Makhija et al., 2016). Given the altered physical connectivity between cytoskeleton and nucleus in DYT1 dystonia patient-derived fibroblasts, we reasoned that these cells may exhibit altered nuclear morphology. To quantify nuclear shape, we measured common metrics including aspect ratio and circularity. We found that nuclei in fibroblasts isolated from DYT1 dystonia patients have a slightly larger aspect ratio than normal fibroblast nuclei, reflecting how they are more elongated as compared to controls (**Figures 4A, B**). Consistent with this, we also found DYT1 dystonia patient-derived fibroblasts have an increased cellular aspect ratio compared to normal control fibroblasts (**Figures 4A, C**). We further investigated nuclear circularity; this shape parameter is sensitive to irregular shapes that deviate from a circle, as

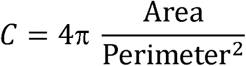

and *C* = 1 for a perfect circle. However, we observed only minor differences in nuclear circularity between DYT1 dystonia patient-derived and normal fibroblasts (**Figure 4D**), which is consistent with the slightly elongated nuclear shapes that we observed in the fibroblasts isolated from DYT1 dystonia patients. Our findings that the DYT1 dystonia-causing ΔE mutation does not have a major impact on nuclear shape contrasts the known effects of reductions or mutations in A-type lamins (Rowat et al., 2013; Harada et al., 2014; Reis-Sobreiro et al., 2018), which are associated with nuclear blebbing, or lobulations, and tend to markedly reduce nuclear circularity (Funkhouser et al., 2013; Rowat et al., 2013; Reis-Sobreiro et al., 2018).

**Figure 4.**
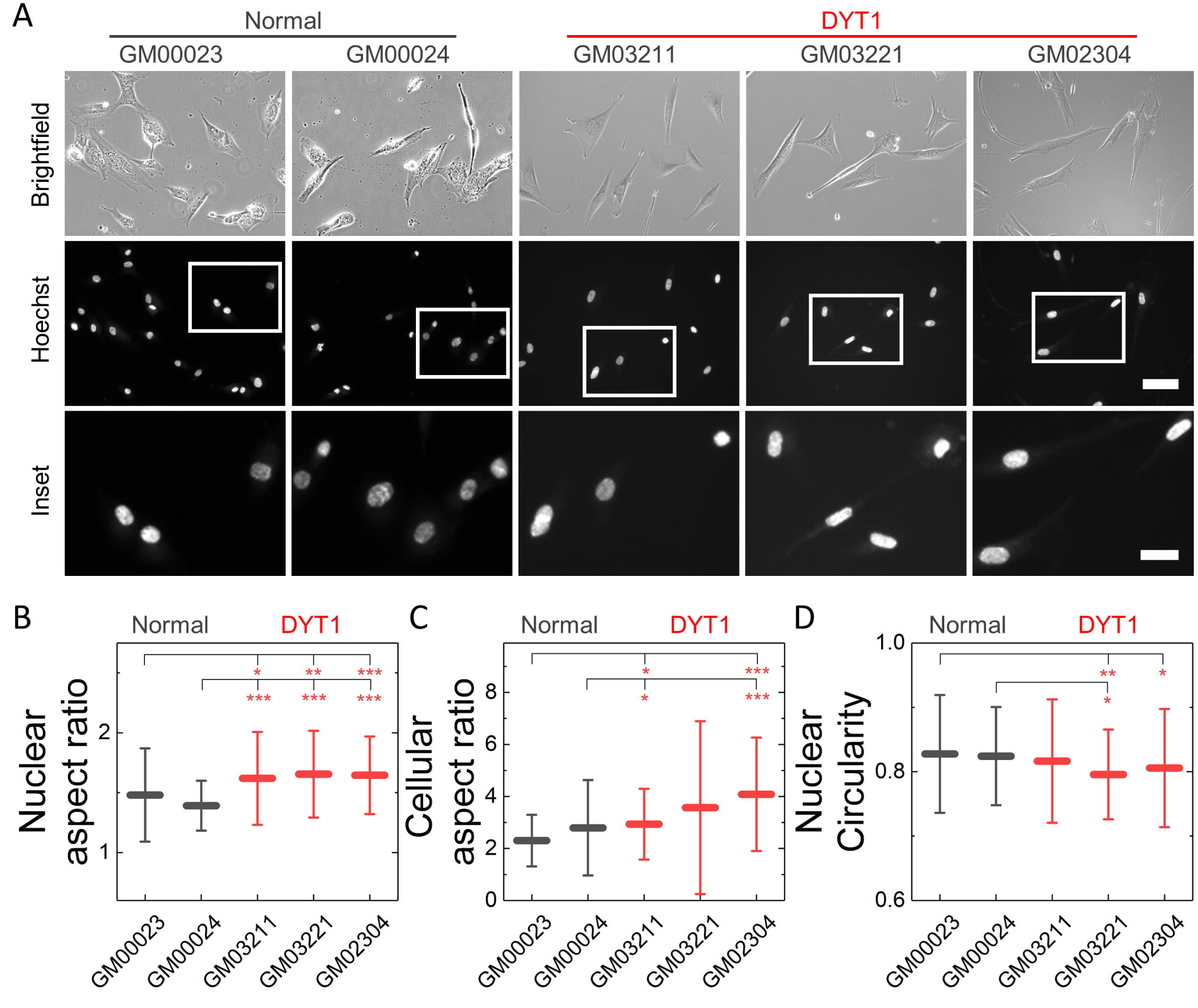
DYT1 patient-derived fibroblasts and nuclei are more elongated. (**A**) Representative brightfield and epifluorescence images of patient-derived normal and DYT1 fibroblasts (brightfield) and nuclei (Hoechst). Scale, 20 µm. Inset: Scale, 10 µm. Quantification of (**B**) nuclear aspect ratio, (**C**) cellular aspect ratio, and (**D**) nuclear circularity. Each data point represents mean ± SD. Data obtained from three independent experiments for N > 30 cells. Statistical significance was determined using Mann-Whitney U test. *** p < 0.001; ** p < 0.01; * p < 0.05; not significant (NS) p > 0.05 is not indicated on these plots for clarity.

### 3.5 DYT1 patient fibroblasts are more susceptible to damage following mechanical stretch

Nuclear lamins are critical for cell survival following exposure to physical forces, suggesting that the mechanical stability of the nucleus imparts protection from external mechanical stresses (Dahl et al., 2004; Denais et al., 2016; Raab et al., 2016; Irianto et al., 2017; Chen et al., 2018; Kim et al., 2018). Since fibroblasts isolated from DYT1 dystonia patients are more deformable than controls, we next tested the hypothesis that DYT1 fibroblasts exhibit increased cell death following exposure to mechanical forces. To test this hypothesis, we grew DYT1 dystonia patient-derived fibroblasts (GM03211, GM02304) and normal controls (GM00023, GM00024) on an elastic collagen-coated PDMS membrane and subjected the resultant membrane with adhered cells to uniaxial mechanical stretch (5% strain). To quantify the magnitude of strain experienced by nuclei, we acquired images of cells in static and stretched conditions. We found that DYT1 dystonia patient-derived fibroblasts cells exhibited larger changes in nuclear area (strain) compared to control fibroblasts in response to the same magnitude of strain of the underlying substrate (**Figures 5A, B**). These observations of increased nuclear strain are consistent with the DYT1 fibroblast nuclei being more deformable than those in control fibroblasts. These findings also suggest that DYT1 dystonia fibroblast nuclei are deforming more in response to external mechanical stresses than control fibroblast nuclei despite the requirement of torsinA for nuclear-cytoskeletal coupling via the LINC complex (Saunders et al., 2017).

**Figure 5.**
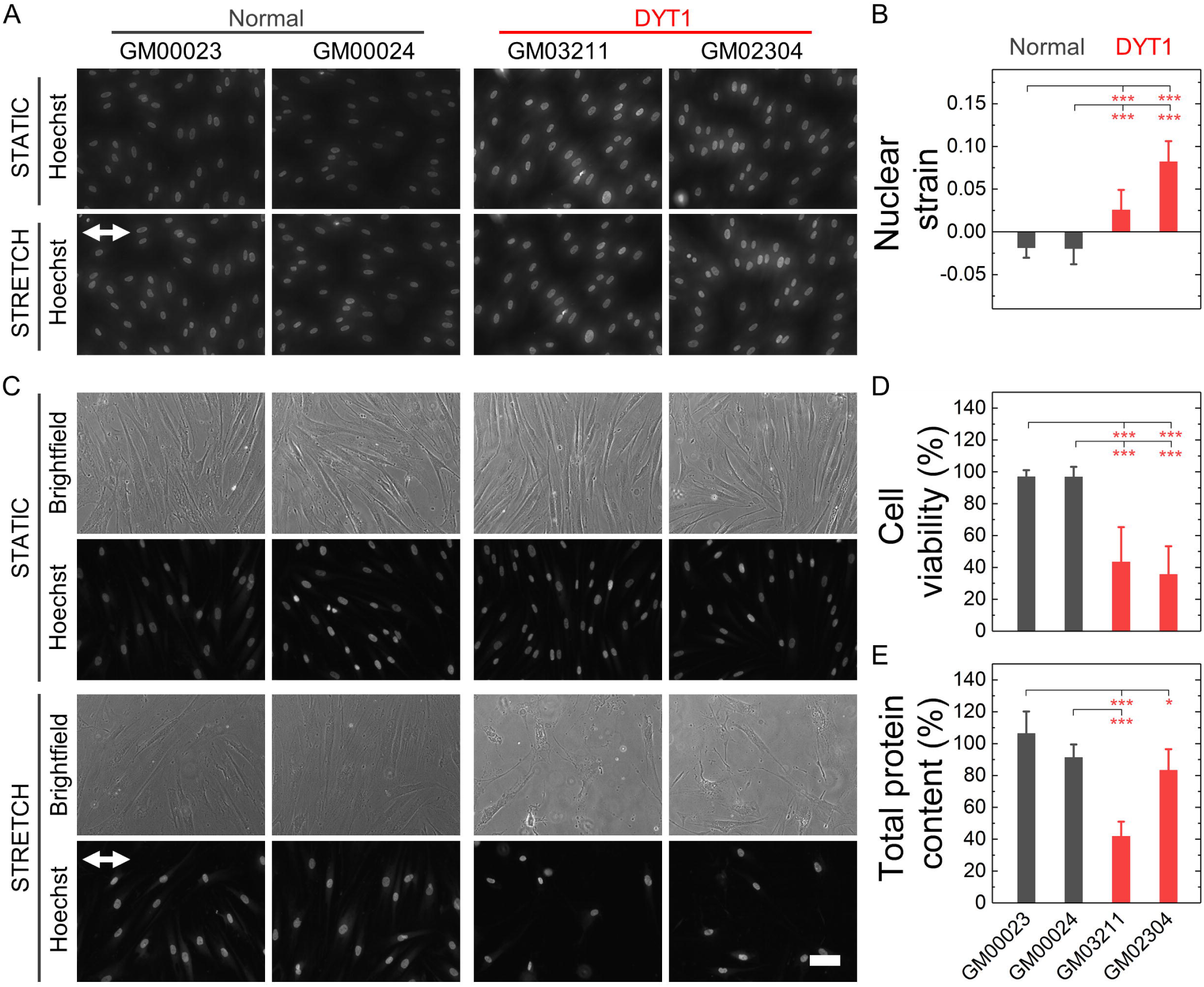
DYT1 patient-derived fibroblasts exhibit increased nuclear strain and are more susceptible to damage upon mechanical stretch. (**A**) Representative images of patient-derived normal and DYT1 fibroblasts stained with the nuclear dye Hoechst. Nuclear morphology was examined after cells were exposed to mechanical stretch (5% strain) for 5 min. Images show stretched (STRETCH) or non-stretched (STATIC) conditions. White arrow denotes direction of uniaxial stretch. (**B**) Quantification of nuclear strain (change in nuclear area) due to cell stretching. N > 15 cells. Statistical significance was determined using Mann-Whitney U test. (**C**) Representative images of patient-derived normal and DYT1 fibroblasts after exposure to 24 h cyclical stretch (5% strain) and static conditions. Cells were stained with Hoechst to visualize nuclei by fluorescence microscopy. Scale, 20 µm. (**D**) Cell viability of normal and DYT1 fibroblasts after exposure to stretch and static conditions. The viability data of stretched cells is normalized to static control for each cell line. N > 500 cells. (**E**) The percentage of cells that are adhered to the PDMS membranes after stretch is quantified by total protein content. Total protein content of stretched cells normalized to static control for each cell line. Each data point represents mean ± SD. Data obtained from two independent experiments. Scale, 20 µm. Statistical significance was determined using Student’s t-test. *** p < 0.001; ** p < 0.01; * p < 0.05; and not significant (NS) p > 0.05 is not indicated on these plots for clarity.

Since the mechanical stability of the nucleus is critical for cell survival following exposure to mechanical stresses (Harada et al., 2014; Denais et al., 2016; Raab et al., 2016; Irianto et al., 2017), we next determined the viability of DYT1 dystonia patient-derived and control fibroblasts after repeated cycles of stretch and relaxation at 5% strain and 0.5 Hz over 24 h. Visual inspection of fibroblasts in stretched versus static samples revealed major morphological differences between fibroblasts isolated from DYT1 dystonia patients compared to control cells after 24 h. In contrast to the aligned morphologies of control fibroblasts, which appear similar in both stretched and static samples, the stretched DYT1 fibroblasts were misaligned and exhibited irregular shapes (**Figure 5C**). To evaluate their viability after stretching, we acquired images of these cells stained with propidium iodide and used quantitative image analysis to count the number of dead cells in each sample. While normal fibroblasts showed no significant cell death after stretching, we observed a marked 57-62% reduction in the viability of DYT1 fibroblasts compared to static control, indicating their reduced survival following exposure to mechanical stresses (**Figure 5D, Supplementary Figure 2**). Since the response of cells to stretch depends on cell-substrate adhesions, we also assessed the number of cells that remained adhered to the PDMS substrate after stretching by quantifying the total protein content of cells lysed from the PDMS membrane. We found a significant ∼17-58% reduction in protein content for the stretched DYT1 cells relative to static control, showing that there was significant detachment of cells from the substrate over the 24 h stretching period. By contrast, over 90% of the control cells were adhered to the membrane after 24 h (**Figure 5E**). The increased detachment of DYT1 dystonia patient-derived fibroblasts is consistent with a previous report of altered integrin-mediated adhesion in these cells (Hewett et al., 2006; Kim et al., 2018). Collectively, these findings indicate a striking difference in the response of DYT1 dystonia patient-derived fibroblasts to mechanical stretch, which could be attributed to their reduced mechanical stability, altered mechanosensation, and/or impaired mechanosignaling. As molecular mediators of mechanotype are generally conserved across cell types (Chang et al., 2015a), we anticipate that our observations of altered mechanotype in human and mouse fibroblasts may also be observed in neurons, which may have important consequences for DYT1 dystonia pathogenesis.

## 4 Discussion

Here we show that DYT1 dystonia is characterized by defective cellular mechanobiology. We find that fibroblasts derived from DYT1 dystonia patients have a more compliant mechanotype than control fibroblasts; these findings are substantiated by the increased deformability of fibroblasts that are torsinA-null; that express torsinA ΔE; or that lack functional LINC complexes. It is intriguing to speculate how altered cellular mechanotype may impact DYT1 dystonia pathogenesis. Mechanotype is closely linked to cellular functions that involve physical force generation, such as motility, and mechanosensing (Anselme et al., 2018; Prahl and Odde, 2018), both of which are required during tissue morphogenesis where cells generate physical forces and sense mechanical cues (Heisenberg and Bellaiche, 2013; Pizzolo et al., 2017). Previous studies showed impaired migration of neurons in the dorsal forebrain of *Tor1a*^-/-^ mouse embryos in vivo (McCarthy et al., 2012) and reduced motility of torsinA-null MEFs and DYT1 dystonia patient-derived fibroblasts in vitro (Nery et al., 2008; Nery et al., 2014). Our observations of the reduced survival of DYT1 dystonia-derived fibroblasts following mechanical stretch may also suggest that these cells have impaired mechanosensation. Like all cells, neurons adapt their own mechanotype by translating mechanical stimuli from their environment into biochemical signals through a process known as mechanotransduction (Tyler, 2012; Franze, 2013). During development, the brain exhibits evolving stiffness gradients due to the variations in the composition and architecture of the extracellular matrix (Franze, 2013; Barnes et al., 2017), which provide mechanical signals that instruct neuronal differentiation, proliferation, and survival (Iwashita et al., 2014; Koser et al., 2016). It is interesting to note that the brains of *Tor1a*^-/-^ mice exhibit elevated LINC complex levels in the proliferative zone and an abnormal morphogenesis characterized by excess neural tissue (Dominguez Gonzalez et al., 2018). Since the brain morphogenesis phenotype observed in the *Tor1a*^-/-^ mice was rescued by the SUN2-deletion (Dominguez Gonzalez et al., 2018), LINC complex regulation of mechanotype may be an important consequence of torsinA activity during brain development in DYT1 dystonia. Taken together with the findings presented here, these results support the hypothesis that DYT1 dystonia may be characterized by defective mechanobiology.

Having established here that torsinA regulates cellular mechanotype and resistance to damage caused by mechanical forces, future efforts will focus on defining the underlying molecular mechanisms. Our findings are consistent with a model where torsinA, LINC complexes, and LINC complex-associated proteins each contribute to cellular mechanotype. Future molecular-level mechanotyping or imaging studies will define the extent to which protein-protein interactions between these structural components versus the individual proteins contributes to cellular and nuclear mechanical stability. Since many nuclear envelope proteins also interact with chromatin and regulate genomic organization (Van de Vosse et al., 2011; Zuleger et al., 2011), changes in gene expression may further impact mechanotype in DYT1 dystonia. Chromatin organization is also a determinant of cellular mechanotype (Pajerowski et al., 2007; Chalut et al., 2012; Schreiner et al., 2015; Stephens et al., 2017). Moreover, torsinA is implicated in other fundamental cellular functions including lipid metabolism and nuclear-cytoplasmic transport (Saunders and Luxton, 2016; Cascalho et al., 2017; Chase et al., 2017), which may also contribute to mechanotype. Identifying the elusive substrate(s) remodeled by torsinA within the contiguous lumen of the endoplasmic reticulum and nuclear envelope should provide further insight into the relative contribution of these torsinA-dependent processes to cellular mechanotype.

While torsinA is required for functional LINC complexes, which physically connect the nucleus and cytoskeleton (Chang et al., 2015a), it is interesting to note that DYT1 dystonia patient-derived fibroblasts exhibit increased nuclear strain following mechanical stretch, suggesting that there are still physical forces pulling on the nucleus during stretch of the underlying substrate. The increased nuclear strain may be explained by force transmission that is mediated by other protein-protein interactions; for example, transmission of external forces to the nucleus could be regulated by microtubules (Alam et al., 2014; Chang et al., 2015a) and/or intermediate filaments, although the relationship between intermediate filaments and DYT1-causing torsinA mutations remains to be fully defined (Hewett et al., 2006; Nery et al., 2014; Saunders et al., 2017). The increased nuclear strain observed in DYT1 fibroblasts is consistent with the loss of torsinA function making the nuclear envelope more deformable, thus resulting in a larger increase in nuclear area for the same magnitude of substrate stretch. We additionally discovered that the altered mechanotype of DYT1 patient fibroblasts is associated with decreased survival following mechanical stretch. These findings are consistent with previous reports that reduced levels of lamin A/C in fibroblasts result in increased cell death following the migration of cells through narrow gaps (Wang et al., 2018). The decreased viability of DYT1 dystonia patient-derived fibroblasts could also result from increased nuclear rupture and double stranded DNA breaks that have been observed in fibroblasts as well as cancer and immune cells (Lammerding et al., 2005; Raab et al., 2016; Isermann and Lammerding, 2017); this resultant damage from mechanical stresses depends on lamin A/C expression levels, suggesting that the mechanical stability of the nuclear envelope is critical for cell survival. Our discovery of the reduced viability of DYT1 fibroblasts following mechanical stretch could also be explained by the altered apoptotic signaling that is triggered by mechanical stimuli (Raab et al., 2016). Future studies will investigate the mechanism of altered survival of DYT1 patient fibroblasts in more detail.

Finally, it is intriguing to speculate that defective mechanobiology may be a common cellular phenotype across different forms of dystonia. The shared disease mechanisms underlying the 25 other known inherited forms of dystonias remain poorly understood, but some evidence suggests altered mechanobiology may be implicated. For example, DYT2 dystonia is caused by an autosomal recessive mutation in the *HPCA* gene, which encodes the calcium-binding hippocalcin protein (Charlesworth et al., 2015) that is implicated in the regulation of the mechanoactive extracellular signal-regulated kinase-mediated signal transduction (Huang and Ingber, 2005; Braunewell et al., 2009). Other known mutations that give rise to dystonia occur in proteins known to play a role in cellular/tissue mechanotype, including β4-tubulin, collagen-6A3, and sarcoglycan-ε (Verbeek and Gasser, 2016). For example, mutations in THAP1 (DYT6) and sarcoglycan-ε (DYT11) cause DYT6 and DYT11 dystonias; these proteins both interact with torsinA (Ledoux et al., 2013). Interestingly, torsinA and sarcoglycan-ε function together to promote proper neurological control over movement (Yokoi et al., 2010). Future studies will determine how broadly altered cellular mechanotype is conserved across different forms of dystonia. Such investigations could also shed light on how TBI can trigger dystonia symptoms, even for individuals that do not carry a known genetic mutation but may be predisposed to acquiring TBI-induced dystonia due to a mutation in another mechanoregulating gene. The motility of neurons is especially relevant in dystonia pathogenesis following TBI, where the directed migration of neurons and neural stem cells is essential for regeneration and repair (Ibrahim et al., 2016). The external mechanical stresses of TBI also result in cell death, which can have consequences for dystonia pathogenesis (Silver and Lux, 1994; Raghupathi, 2004). A deeper understanding of the mechanobiology of dystonia could further drive the discovery of novel therapeutic targets for treatment of this debilitating disease.

## Supporting information

## 5 Abbreviations

AAA+: ATPase associated with various cellular activities
*d*: Cell diameter
*E*_*a*_: Apparent cell elastic modulus
*C*: Circularity
LAP1: Lamina-associated polypeptide 1
LULL1: Luminal domain-like LAP1
LINC: Linkers of nucleoskeleton and cytoskeleton
MEF: Mouse embryonic fibroblast
PMF: Parallel microfiltration
q-DC: Quantitative deformability cytometry
SUN: Sad1/UNC-84
TAN: Transmembrane actin-associated nuclear
*TT*: Transit time
WT: Wild type

## 7 Author Contributions

NKG, ACR and GWGL designed the research. NKG performed all experiments and analyzed data. CL performed q-DC experiments. PK and NKG conducted cell stretching experiments. CAS, LGF, and SGY generated MEF cell lines. NKG, ACR and GWGL wrote the manuscript. All authors reviewed the manuscript.

## 9 Acknowledgments

We thank members of the Luxton, Rowat, and Young laboratories for helpful discussions. Special acknowledgments go to W. T. Dauer for the *Tor1a* and *Tor1aip1* MEFs as well as B. Burke for the *SUN2* and *SUN1/2* MEFs.

