## Supplementary material for "DYT1 dystonia patient-derived fibroblasts have increased deformability and susceptibility to damage by mechanical forces"

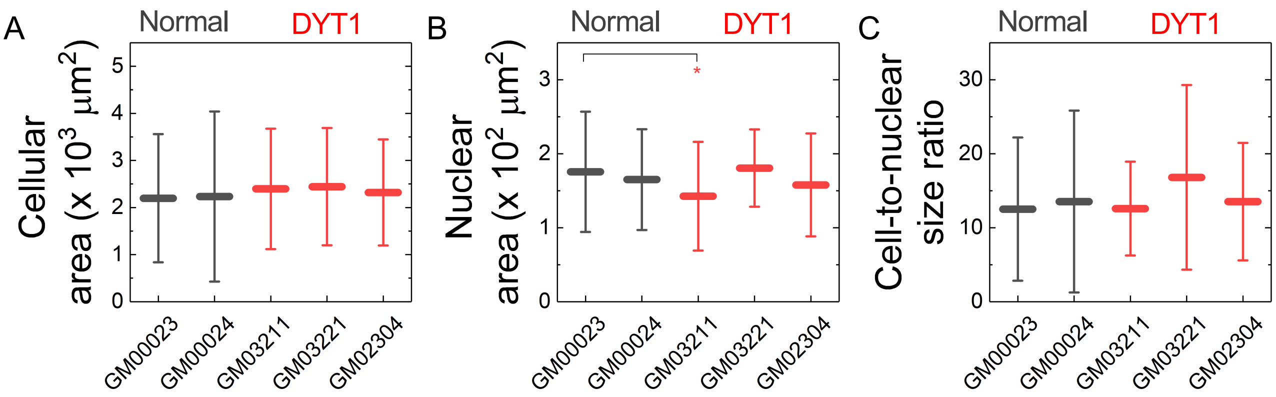


**Supplementary Figure 1. Quantification of cell and nuclear area of normal and DYT1 fibroblasts.** (**A**) Cellular area, (**B**) nuclear area and, (**C**) cell-to-nucleus area ratio for indicated patient-derived normal and DYT1 fibroblasts. Each data point represents mean ± SD. Data obtained from three independent experiments. Statistical significance was determined using Mann-Whitney U test and indicated where significant. * $p < 0.05$. Not significant (NS) $p > 0.05$ is not indicated on these plots for clarity.

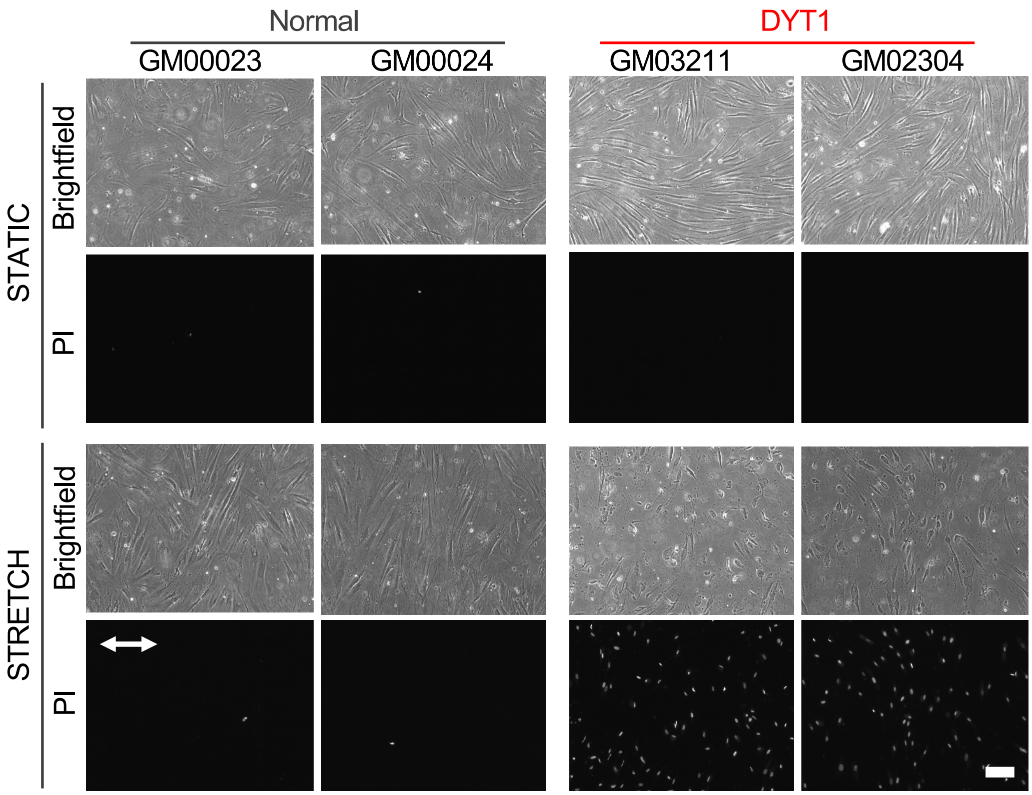


**Supplementary Figure 2. DYT1 fibroblasts have reduced viability after mechanical stretching compared to normal fibroblasts.** Representative images of PI-labeled cells, which indicate cell death, after stretching (stretch) compared to without mechanical stretching (static). White arrow shows the direction of uniaxial stretch. Scale, 100 µm.
